# Spontaneous eye blink rate and dopamine synthesis capacity: Preliminary evidence for an absence of positive correlation

**DOI:** 10.1101/215178

**Authors:** Guillaume Sescousse, Romain Ligneul, Ruth J. van Holst, Lieneke K. Janssen, Femke de Boer, Marcel Janssen, Anne S. Berry, William J. Jagust, Roshan Cools

## Abstract

Dopamine is central to a number of cognitive functions and brain disorders. Given the cost of neurochemical imaging in humans, behavioral proxy measures of dopamine have gained in popularity in the past decade, such as spontaneous eye blink rate (sEBR). Increased sEBR is commonly associated with increased dopamine function based on pharmacological evidence and patient studies. Yet, this hypothesis has not been validated using *in vivo* measures of dopamine function in humans. In order to fill this gap, we measured sEBR and striatal dopamine synthesis capacity using [^18^F]DOPA PET in 20 participants (9 healthy individuals and 11 pathological gamblers). Our results, based on frequentist and Bayesian statistics, as well as region-of-interest and voxel-wise analyses, argue against a positive relationship between sEBR and striatal dopamine synthesis capacity. They show that, if anything, the evidence is in favor of a negative relationship. These results, which complement findings from a recent study that failed to observe a relationship between sEBR and dopamine D2 receptor availability, suggest that caution and nuance are warranted when interpreting sEBR in terms of a proxy measure of striatal dopamine.

## Introduction

Dopamine is a key ascending neuromodulator subserving a range of cognitive functions, including motivation, learning and attention. In turn, striatal dopamine dysregulation is a common feature across many psychiatric disorders in which those functions are impaired, including addiction, schizophrenia, attention deficit hyperactivity disorder, and depression (Grace, 2016; Nutt et al., 2015; Russo and Nestler, 2013). Unfortunately, direct assessment of dopamine function in humans is only possible with expensive and invasive techniques such as positron emission tomography (PET), which require complex infrastructure and in-depth methodological expertise. As a consequence, non-invasive proxy measures have gained in popularity. In particular, spontaneous eye-blink rate (sEBR) has been used in many studies to investigate dopamine’s role in various cognitive tasks and patient populations (for a comprehensive review see Jongkees and Colzato, 2016).

Evidence supporting a link between dopamine function and sEBR comes primarily from pharmacological studies in both humans and animals. These studies have shown that dopamine-enhancing drugs generally increase sEBR (Karson, 1983; Strakowski et al., 1996), whereas dopamine-decreasing drugs tend to decrease sEBR (Lawrence and Redmond, 1991; Mavridis et al., 1991) – although a number of other studies have reported null or opposite results (Ebert et al., 1996; Kleven and Koek, 1996; Kotani et al., 2016; Mohr et al., 2005; van der Post et al., 2004). Similarly, brain disorders characterized by a hypo-dopaminergic state, such as Parkinson’s disease, have been mostly associated with decreased sEBR (Chen et al., 2003; Deuschl and Goddemeier, 1998), whereas disorders characterized by a hyper-dopaminergic state, such as schizophrenia, have been associated with increased sEBR (Chan et al., 2010; Chen et al., 1996). These observations have led to the pervasive hypothesis that sEBR is *positively* correlated with dopamine function (Jongkees and Colzato, 2016).

However, establishing sEBR as a reliable marker of human dopamine function requires direct evidence for a relationship between sEBR and *in vivo* measures of dopamine function in humans. Dopamine function can be measured in several ways, for example, in terms of synthesis capacity, transporter density and/or (changes in) receptor availability. Only two PET studies to date have tackled this question and both of these focused on dopamine receptor availability. The first study by Groman et al. (2014) measured dopamine D2 and D1 receptor availability in vervet monkeys, using [^18^F]fallypride and [^11^C]NNC-112 PET, respectively. The results showed that striatal dopamine D2 (but not D1) receptor availability was positively related with sEBR, as well as with D2 receptor agonist-induced changes in sEBR. However, a recent study by Dang et al. (2017) was unable to replicate these results in humans. Using [^18^F]fallypride PET and the dopamine D2 receptor agonist bromocriptine, they found that dopamine D2 receptor availability was neither related to baseline sEBR or bromocriptine-induced changes in sEBR. In the present study we focused on another aspect of dopamine function, namely striatal dopamine synthesis capacity, as measured by [^18^F]DOPA PET. Based on the above described literature and the commonly assumed positive relationship between sEBR and dopamine function (Jongkees and Colzato, 2016), we hypothesized that dopamine synthesis capacity would be positively related with sEBR. This hypothesis is further in line with a previous report showing that sEBR is positively correlated with post-mortem measures of dopamine concentration in the caudate nucleus of monkeys treated with the neurotoxic drug MPTP (Taylor et al., 1999).

## Experimental procedures

### Participants

Thirty male participants were recruited, among which were 15 pathological gamblers (PGs) and 15 matched healthy controls (HCs). The PET data from these participants have been reported in a previous study comparing baseline dopamine synthesis capacity between groups (van Holst et al., 2017). PGs were recruited through advertisement and addiction treatment centers, and reported not to be medicated or in treatment for their gambling at the time of the PET study. HCs were recruited through advertisement. All gamblers qualified as PGs as they met ≥5 DSM-IV-TR criteria for pathological gambling. Current drug use disorder was a reason for exclusion for all participants. Additionally, participants were excluded if they were currently following psychiatric treatment, drank more than four alcoholic beverages daily, used marijuana more than once per month, were using (psychotropic) medication, had a lifetime history (as assessed with the MINI interview) of schizophrenia, bipolar disorder, attention deficit hyperactivity disorder, autism, bulimia, anorexia, anxiety disorder, obsessive compulsive disorder, or had a past 6-month history of major depressive episode.

Nine participants (3 PGs; 6 HCs) were not included due to technical problems with the electro-oculography (EOG) recordings (data loss > 50% of the total recording time). Data loss was related to EOG amplifier saturation (output voltage of the amplifier exceeding its operating range sensitivity). Three participants were kept in the analyses despite minor data loss (exploitable data: 3.5, 4.4 and 4.5 min). One gambler was further excluded because he met the DSM-IV-TR criteria for past year marijuana dependence, which is known to influence sEBR measures (Kowal et al., 2011). As a result, the final sample comprised 20 participants (mean age: 38.2, age range: 22-54, gamblers: n=11, controls: n=9).

All participants provided written informed consent to take part in the study, which was approved by the regional research ethics committee (Commissie Mensgebonden Onderzoek, region Arnhem-Nijmegen, registration number NL41522.091.12).

### Study procedures

The study took place at the Radboud University Medical Center. Upon arrival of the participants, spontaneous eye-blink rate (sEBR) was measured using EOG recordings (see detailed description below). Approximately one hour before entering the PET scanner, participants received 150 mg of carbidopa and 400 mg of entacapone to reduce peripheral metabolism of [^18^F]DOPA and increase tracer availability in the brain while having no psychotropic side effects. The participants further performed computerized tasks not reported here.

### Spontaneous eye-blink rate: recording and analyses

We followed standard procedures for the acquisition, preprocessing and analysis of the EOG data (Colzato et al., 2009; Slagter et al., 2010). The data were acquired during the day (3:30-4:30pm) over a 6-min period, using two vertical and two horizontal Ag-AgCl electrodes placed around the left eye. The sampling rate of 100Hz. The vertical EOG signal (vEOG), used for eye-blink assessment, was obtained from a bipolar montage using the electrodes placed above and below the eye. The horizontal EOG (hEOG) signal, used to exclude artifacts produced by saccades, was obtained from a bipolar montage using the electrodes placed lateral to the external canthi.

Participants were comfortably seated and instructed to fixate the wall in front of them while they believed the system was being calibrated and the experimenter was outside of the room. They were not instructed in any way about blinking, and were asked not to move their head or activate their facial muscles in order to minimize EOG artifacts.

The vEOG signals was rectified and band-pass filtered between 0.5 and 20 Hz. Eye blinks were detected using an automated procedure based on a voltage change threshold of 100 µV in a time interval of 400 ms. The vEOG signal was then visually inspected by two of the authors (R.L. and G.S.) in order to assess detection accuracy (i.e. presence/absence of blinks) and remove potential artefacts resulting from muscle activity and saccades as detected in the hEOG signal. Since the inter-rater reliability was very high (Cronbach’s α = 0.989), the resulting sEBR measures from the two raters were averaged for subsequent analyses.

### PET and MRI data acquisition

All PET scans were acquired at the department of Nuclear Medicine of the Radboud University Medical Center using a Siemens mCT PET/CT camera (with 40 slice CT, voxel size 4x4 mm in-plane, 5mm slice thickness). Participants were positioned as comfortably as possible, in a supine position, with the head slightly fixated in a headrest to avoid movement. First a low dose CT was made for attenuation correction of the PET images, followed by an 89-minute dynamic PET scan. The scan started at the same time as the bolus injection of the [^18^F]DOPA into an antecubital vein. Images were reconstructed with an ordered subset expectation maximization algorithm with weighted attenuation, time-of-flight recovery, scatter corrected, and smoothed with a 4-mm FWHM kernel.

A high-resolution anatomical scan (T1-weighted MP-RAGE, TR = 2300 ms, TE = 3.03 ms, 8° flip-angle, 192 sagittal slices, slice-matrix size= 256x256, voxel size 1x1x1 mm3) was obtained using a 3 Tesla MR Siemens scanner at the Donders Centre for Cognitive Neuroimaging and used for co-registration with the PET data.

### PET data preprocessing and analyses

We realigned the [^18^F]DOPA images to the middle (11^th^) frame to correct for movement during scanning using SPM8 (http://www.fil.ion.ucl.ac.uk/spm/). The mean [^18^F]DOPA image and the realigned frames were coregistered to the structural MRI scan using SPM8. Higher [^18^F]DOPA uptake, indexed by higher Ki values, is established to reflect higher dopamine synthesis capacity (Snow et al., 1993). To create Ki images representing the amount of tracer accumulated in the brain *relative* to a cerebellar region of reference, we used an in-house graphical analysis program implementing Patlak plotting (Patlak and Blasberg, 1985). Ki images were generated from PET frames corresponding to 24 to 89 minutes. These images are comparable to Ki images obtained using a blood input function but are scaled to the volume of tracer distribution in the reference region. Finally, structural MNI scans were normalized to a standard MNI template (IBASPM152) and the transformation matrix was applied to coregistered Ki images. As expected from previous studies (Rakshi et al., 1999), the voxels with the highest Ki values were located in the striatum and the brainstem (Figure 1A).

**Figure 1.**
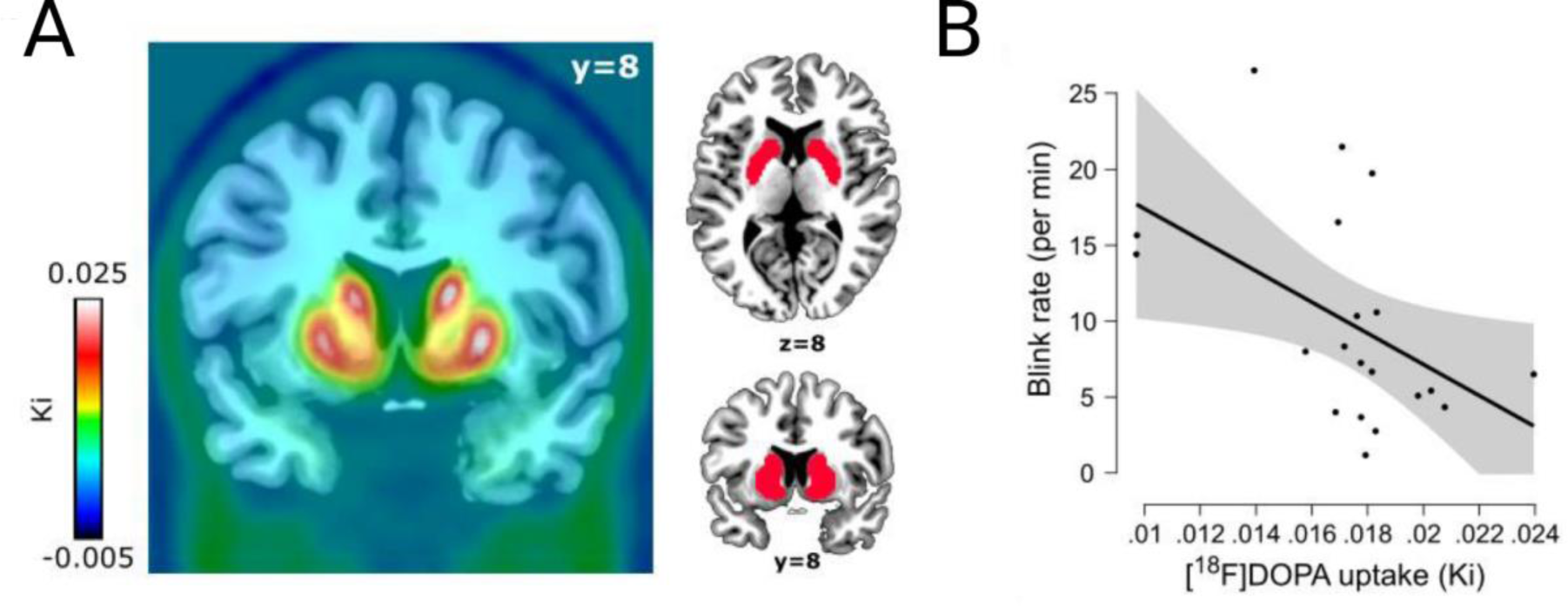
Main results: mapping of dopamine synthesis capacity and relationship with spontaneous eye blink rate in the striatum. **(A)** On the left, whole-brain map of mean Ki values across the twenty participants included in the analyses. On the right, the mask used for the region-of-interest analyses is depicted in red. **(B)** Negative correlation between spontaneous eye blink rate and Ki values in the striatal mask. The shaded area represents the 95% confidence interval.

For the main analyses, cerebellum-corrected Ki values were extracted from a striatal region of interest defined independently from an [^18^F]DOPA PET template normalized to MNI space (García-Gómez et al., 2013). In order to improve sensitivity, we retained the template voxels in which Ki values exceeded the mean Ki value across all voxels by at least 3 standard deviations (Figure 1A).

For the exploratory voxel-wise analysis (Figure 2B), we used SPM8 to perform a linear regression in which global signal intensity was included as a covariate of no-interest. The analysis was restricted to an anatomical mask covering the entire striatum (Choi et al., 2012), within which the results were corrected for multiple comparisons using a family-wise error (FWE) corrected threshold of p<0.05.

**Figure 2.**
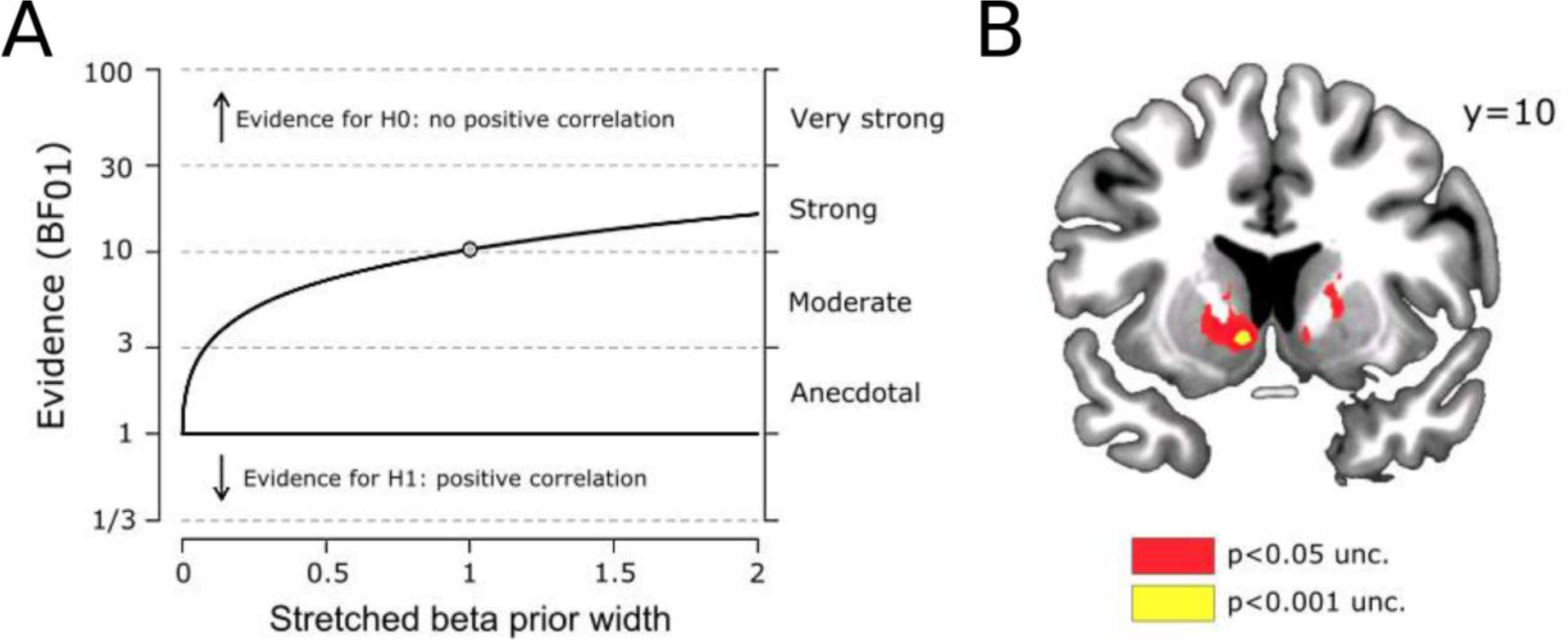
Control and exploratory analyses. **(A)** Robustness check for the Bayesian correlation analysis. This graph shows that the Bayes factor quantifying the relative evidence for the absence vs presence of a positive correlation exceeds the critical threshold of 3 for a large range of beta prior widths, even extending to strong prior beliefs in the existence of a positive correlation (values < 1 correspond to a prior biased in favor of a positive correlation, values > 1 correspond to a prior biased in favor of an absence of a positive correlation, and value = 1 corresponds to an uninformative (flat) prior as used in our main analysis). **(B)** A voxel-wise analysis restricted to an anatomical mask of the striatum shows that the negative correlation between sEBR and Ki values peaks in the left nucleus accumbens ([x y z] = [−8 10 −6]; p_FWE striatal mask_<0.05).

### Statistics

Since extracted Ki values were non-normally distributed (Kolmogorov-Smirnov test: p<0.003), we used rank correlation coefficients (Spearman ρ). We also performed partial regression analyses, in which beta coefficients were estimated using bootstrapping in SPSS. Bayesian analyses were conducted using JASP (Wagenmakers et al., 2017).

## Results

In contrast to our prediction, frequentist statistics indicated a negative relationship between sEBR and Ki values (Spearman ρ = −0.504, p=0.024, Figure 1B). As both age and gambling status are known to be related to Ki values (Berry et al., 2016; van Holst et al., 2017), we next regressed sEBR onto Ki values while adding these two variables as covariates, revealing a marginally significant negative relationship between sEBR and dopamine synthesis capacity (beta=-1.82, p=0.09).

Given the unexpectedness of this finding, we sought to quantify the evidence for the null hypothesis that sEBR and Ki values are *not positively* correlated (H0), as opposed to *positively* correlated (H1). We used a Bayesian analysis with an uninformative (flat) prior distribution over the (0,1) interval. The relative evidence in favor of H0 over H1 was strong (Bayes factor: BF_01_=10.34), indicating that our data are ten times more likely under the null hypothesis of *no positive relationship* than under the alternative hypothesis of *a positive relationship*. Importantly, a Bayes factor robustness check showed that this conclusion holds even when using strong prior beliefs in the existence of a positive correlation (Figure 2A). In fact, there was also moderate evidence (BF_10_=4.90) in favor of a negative relationship between sEBR and Ki values (H1 versus H0 defined as an absence of negative relationship).

Finally, an exploratory voxel-wise analysis showed that the negative relationship between sEBR and Ki values was strongest in the left nucleus accumbens (Figure 2B).

### Control analyses

In order to assess the robustness of our results, we performed a number of control analyses. First, we re-ran our analyses after excluding the data from the three participants with incomplete sEBR datasets. The results remained unchanged, showing a negative correlation between sEBR and striatal dopamine synthesis (Spearman ρ=-0.5, p=0.016), while the Bayesian correlation analysis indicated moderate to strong evidence against hypothesis H1 of a positive correlation (BF_01_=9.95). When splitting the data as a function of gambling status, we observed a negative association between sEBR and Ki values in both groups (healthy controls (n=9): ρ=-0.63, p=0.067; pathological gamblers (n=11): ρ=-0.46, p=0.15). Although p-values did not reach significance, Bayesian analyses similarly revealed moderate evidence against a positive correlation (healthy controls: BF_01_=5.66; pathological gamblers: BF_01_=6.29), although of course these values should be interpreted with caution given the small sample sizes of these subgroups.

Finally, we verified that our results were not influenced by the method used to define eye blinks. Indeed, in our main analysis we used a relatively low detection threshold of 100µV, which has the advantage of capturing most blinks (high sensitivity), but the disadvantage of increasing the likelihood of false positives due to muscular artefacts (decreased specificity), which then requires manual (i.e. subjective) editing. To circumvent that issue we re-ran our analyses using more conservative thresholds (200µV and 300µV), which provide increased specificity and thus eliminate the need for manual editing (at the cost of decreased sensitivity). The resulting sEBR measures were again negative correlated with Ki values, although not significantly (200µV: Spearman ρ=-0.41, p=0.077; 300µV: Spearman ρ=-0.38, p=0.099), while corresponding Bayes factors again provided strong evidence in favor of the null hypothesis H0 (200µV: BF_01_=10.23; 300µV: BF_01_=10.29).

## Discussion

Spontaneous EBR has become increasingly popular as a putative behavioral marker of endogenous dopamine. Interestingly, most of the past studies that have used sEBR in this capacity have loosely referred to “dopamine function” or “dopamine activity”, perhaps reflecting the current dearth of knowledge regarding which specific aspect(s) of the dopamine system correlate(s) with sEBR. Here we tested the hypothesis of a positive relationship between sEBR and striatal dopamine synthesis capacity, based on the proposal that sEBR is positively related with striatal dopamine function (Jongkees and Colzato, 2016), and previous results showing a positive correlation between sEBR and striatal dopamine levels measured in post-mortem monkey brains (Taylor et al., 1999). Both frequentist and Bayesian statistics, as well as ROI and voxel-wise analyses, argue against the existence of such a positive relationship in our sample, and show that, if anything, the evidence is in favor of a negative relationship. While we prefer to refrain from making speculative interpretations regarding the existence of a negative relationship, given the moderate level of evidence, we believe these data provide a strong cautionary message against the use of sEBR as a positive predictor of pre-synaptic striatal dopamine.

Importantly, our data speak specifically to dopamine synthesis capacity, and do not preclude correlations of sEBR with other aspects of the dopamine system, including (changes in) dopamine D2 receptor availability. It has been suggested that sEBR might primarily reflect striatal D2 receptor activity, based on the observed positive correlation with D2 receptor availability in monkeys (Groman et al., 2014), and the observation that sEBR better predicts D2-mediated punishment learning than D1-mediated reward learning (Cavanagh et al., 2014; Slagter et al., 2015). However, this relationship between sEBR and dopamine D2 functioning has been recently questioned by a PET study which failed to replicate it in humans (Dang et al., 2017). Also, such a relationship remains difficult to reconcile with recent findings showing a positive association between dopamine D2 receptor availability and dopamine synthesis capacity (Berry et al., 2017, but see Ito et al., 2011), as this would predict a positive relationship between sEBR and dopamine synthesis capacity, in contrast to our results. These inconsistencies certainly call for further research to better elucidate the dopaminergic mechanisms underlying sEBR.

This study is not devoid of limitations. First, our sample size (n = 20) is relatively small, although one should note that it is larger than the samples used in many of the psychopharmacological studies that have investigated dopaminergic drug effects on sEBR (Jongkees and Colzato, 2016). In fact, a power analysis performed with GPower3.1 (Faul et al., 2009) suggests that a sample size of 12 individuals should have been sufficient to replicate with 95% power the positive relationship reported by Taylor et al. (1999) between striatal dopamine levels and sEBR in monkeys (original effect size: R^2^ = 0.62). In addition, as argued by Dang et al. (2017), the use of sEBR as a reliable predictor of dopamine function implicitly requires that the positive relationship between these two variables should be strong and thus observable even in small samples. For these reasons, we believe that the preliminary evidence reported here is valuable, even though a replication in a larger sample size is warranted.

Another aspect that may be perceived as a limitation is the use of a mixed population of healthy participants and pathological gamblers. While we acknowledge that pathological gamblers are not typical individuals and are characterized, among other things, by elevated striatal dopamine synthesis (van Holst et al., 2017), we believe that this is not necessarily an issue in the context of the current study. Indeed, our goal was to examine whether individual differences in sEBR and dopamine synthesis were positively related, regardless of the origin of these individual differences. If sEBR is to be used as proxy measure of dopamine levels, it should be insensitive to the underlying causes of individual variations, so that it can be effectively used in both clinical and non-clinical populations. In fact, a large portion of the literature that has led to the hypothesis of a link between sEBR and dopamine function is based on the study of clinical populations characterized by dopamine dysfunctions. Finally, one should note that restricting our analyses to healthy individuals did not affect the results, still showing moderate evidence against a positive correlation.

To conclude, our study does not support the hypothesis of a positive relationship between sEBR and striatal dopamine synthesis, and if anything, provides evidence against it. Even though it is based on a modest sample size and needs to be replicated in a larger sample – which we are currently attempting to do, it warrants caution for future studies that may be tempted to use sEBR as a proxy measure of striatal dopamine synthesis capacity.

## Authors disclosures

### Contributors

GS, RJvH and RC designed the study. RJvH and LKJ collected the data. GS, RL, RJvH, FdB, MJ, ASB and WJJ performed data analysis. GS and RL wrote the first draft of the manuscript. RJvH and RC edited the manuscript.

## References

Berry, A.S., Shah, V.D., Baker, S.L., Vogel, J.W., O'Neil, J.P., Janabi, M., Schwimmer, H.D., Marks, S.M., Jagust, W.J., 2016. Aging affects dopaminergic neural mechanisms of cognitive flexibility. J. Neurosci. doi:10.1523/JNEUROSCI.0626-16.2016

Berry, A.S., Shah, V.D., Furman, D.J., White, R.L., Baker, S.L., O'Neil, J.P., Janabi, M., D'Esposito, M., Jagust, W.J., 2017. Dopamine Synthesis Capacity is Associated with D2/3 Receptor Binding but not Dopamine Release. Neuropsychopharmacology. doi:10.1038/npp.2017.180

Cavanagh, J.F., Masters, S.E., Bath, K., Frank, M.J., 2014. Conflict acts as an implicit cost in reinforcement learning. Nat. Commun. 5, 5394.

Chan, K.K.S., Hui, C.L.M., Lam, M.M.L., Tang, J.Y.M., Wong, G.H.Y., Chan, S.K.W., Chen, E.Y.H., 2010. A three-year prospective study of spontaneous eye-blink rate in first-episode schizophrenia: relationship with relapse and neurocognitive function. East Asian Arch. Psychiatry 20, 174–179.

Chen, E.Y., Lam, L.C., Chen, R.Y., Nguyen, D.G., 1996. Blink rate, neurocognitive impairments, and symptoms in schizophrenia. Biol. Psychiatry 40, 597–603.

Chen, W.H., Chiang, T.J., Hsu, M.C., Liu, J.S., 2003. The validity of eye blink rate in Chinese adults for the diagnosis of Parkinson's disease. Clin. Neurol. Neurosurg. 105, 90–92.

Choi, E.Y., Yeo, B.T.T., Buckner, R.L., 2012. The organization of the human striatum estimated by intrinsic functional connectivity. J. Neurophysiol. 108, 2242–2263.

Colzato, L.S., van den Wildenberg, W.P.M., van Wouwe, N.C., Pannebakker, M.M., Hommel, B., 2009. Dopamine and inhibitory action control: evidence from spontaneous eye blink rates. Exp. Brain Res. 196, 467–474.

Dang, L.C., Samanez-Larkin, G.R., Castrellon, J.J., Perkins, S.F., Cowan, R.L., Newhouse, P.A., Zald, D.H., 2017. Spontaneous Eye Blink Rate (EBR) Is Uncorrelated with Dopamine D2 Receptor Availability and Unmodulated by Dopamine Agonism in Healthy Adults. eNeuro 4. doi:10.1523/ENEURO.0211-17.2017

Deuschl, G., Goddemeier, C., 1998. Spontaneous and reflex activity of facial muscles in dystonia, Parkinson's disease, and in normal subjects. J. Neurol. Neurosurg. Psychiatry 64, 320–324.

Ebert, D., Albert, R., Hammon, G., Strasser, B., May, A., Merz, A., 1996. Eye-Blink Rates and Depression. Neuropsychopharmacology 15, 332–339.

Faul, F., Erdfelder, E., Buchner, A., Lang, A.-G., 2009. Statistical power analyses using G* Power 3.1: Tests for correlation and regression analyses. Behav. Res. Methods 41, 1149–1160.

García-Gómez, F.J., García-Solís, D., Luis-Simón, F.J., Marín-Oyaga, V.A., Carrillo, F., Mir, P., Vázquez-Albertino, R.J., 2013. Elaboration of the SPM template for the standardization of SPECT images with 123I-Ioflupane. Rev. Esp. Med. Nucl. Imagen Mol. 32, 350–356.

Grace, A.A., 2016. Dysregulation of the dopamine system in the pathophysiology of schizophrenia and depression. Nat. Rev. Neurosci. 17, 524.

Groman, S.M., James, A.S., Seu, E., Tran, S., Clark, T.A., Harpster, S.N., Crawford, M., Burtner, J.L., Feiler, K., Roth, R.H., Elsworth, J.D., London, E.D., Jentsch, J.D., 2014. In the blink of an eye: relating positive-feedback sensitivity to striatal dopamine D2-like receptors through blink rate. J. Neurosci. 34, 14443–14454.

Ito, H., Kodaka, F., Takahashi, H., Takano, H., Arakawa, R., Shimada, H., Suhara, T., 2011. Relation between presynaptic and postsynaptic dopaminergic functions measured by positron emission tomography: implication of dopaminergic tone. J Neurosci. 31, 7886–7890.

Jongkees, B.J., Colzato, L.S., 2016. Spontaneous eye blink rate as predictor of dopamine-related cognitive function-A review. Neurosci. Biobehav. Rev. 71, 58–82.

Karson, C.N., 1983. Spontaneous eye-blink rates and dopaminergic systems. Brain 106 (Pt 3), 643– 653.

Kleven, M.S., Koek, W., 1996. Differential effects of direct and indirect dopamine agonists on eye blink rate in cynomolgus monkeys. J. Pharmacol. Exp. Ther. 279, 1211–1219.

Kotani, M., Kiyoshi, A., Murai, T., Nakako, T., Matsumoto, K., Matsumoto, A., Ikejiri, M., Ogi, Y., Ikeda, K., 2016. The dopamine D1 receptor agonist SKF-82958 effectively increases eye blinking count in common marmosets. Behav. Brain Res. 300, 25–30.

Kowal, M.A., Colzato, L.S., Hommel, B., 2011. Decreased spontaneous eye blink rates in chronic cannabis users: evidence for striatal cannabinoid-dopamine interactions. PLoS One 6, e26662.

Lawrence, M.S., Redmond, D.E., Jr, 1991. MPTP lesions and dopaminergic drugs alter eye blink rate in African green monkeys. Pharmacol. Biochem. Behav. 38, 869–874.

Mavridis, M., Degryse, A.-D., Lategan, A.J., Marien, M.R., Colpaert, F.C., 1991. Effects of locus coeruleus lesions on parkinsonian signs, striatal dopamine and substantia nigra cell loss after 1-methyl-4-phenyl-1, 2, 3, 6-tetrahydropyridine in monkeys: a possible role for the locus coeruleus in the progression of Parkinson's disease. Neuroscience 41, 507–523.

Mohr, C., Sándor, P.S., Landis, T., Fathi, M., Brugger, P., 2005. Blinking and schizotypal thinking. J. Psychopharmacol. 19, 513–520.

Nutt, D.J., Lingford-Hughes, A., Erritzoe, D., Stokes, P.R.A., 2015. The dopamine theory of addiction: 40 years of highs and lows. Nat. Rev. Neurosci. 16, 305–312.

Patlak, C.S., Blasberg, R.G., 1985. Graphical evaluation of blood-to-brain transfer constants from multiple-time uptake data. Generalizations. J. Cereb. Blood Flow Metab. 5, 584–590.

Rakshi, J.S., Uema, T., Ito, K., Bailey, D.L., Morrish, P.K., Ashburner, J., Dagher, A., Jenkins, I.H., Friston, K.J., Brooks, D.J., 1999. Frontal, midbrain and striatal dopaminergic function in early and advanced Parkinson's disease A 3D [18F] dopa-PET study. Brain 122, 1637–1650.

Russo, S.J., Nestler, E.J., 2013. The brain reward circuitry in mood disorders. Nat. Rev. Neurosci. 14, 609–625.

Slagter, H.A., Davidson, R.J., Tomer, R., 2010. Eye-blink rate predicts individual differences in pseudoneglect. Neuropsychologia 48, 1265–1268.

Slagter, H.A., Georgopoulou, K., Frank, M.J., 2015. Spontaneous eye blink rate predicts learning from negative, but not positive, outcomes. Neuropsychologia 71, 126–132.

Strakowski, S.M., Sax, K.W., Setters, M.J., Keck, P.E., Jr, 1996. Enhanced response to repeated d-amphetamine challenge: evidence for behavioral sensitization in humans. Biol. Psychiatry 40, 872–880.

Taylor, J.R., Elsworth, J.D., Lawrence, M.S., Sladek, J.R., Jr, Roth, R.H., Redmond, D.E., Jr, 1999. Spontaneous blink rates correlate with dopamine levels in the caudate nucleus of MPTP-treated monkeys. Exp. Neurol. 158, 214–220.

van der Post, J., de Waal, P.P., de Kam, M.L., Cohen, A.F., van Gerven, J.M.A., 2004. No evidence of the usefulness of eye blinking as a marker for central dopaminergic activity. J. Psychopharmacol. 18, 109–114.

van Holst, R.J., Sescousse, G., Janssen, L.K., Janssen, M., Berry, A.S., Jagust, W.J., Cools, R., 2017. Increased Striatal Dopamine Synthesis Capacity in Gambling Addiction. Biol. Psychiatry. doi:10.1016/j.biopsych.2017.06.010

Wagenmakers, E.-J., Love, J., Marsman, M., Jamil, T., Ly, A., Verhagen, J., Selker, R., Gronau, Q.F., Dropmann, D., Boutin, B., Meerhoff, F., Knight, P., Raj, A., van Kesteren, E.-J., van Doorn, J., Šmíra, M., Epskamp, S., Etz, A., Matzke, D., de Jong, T., van den Bergh, D., Sarafoglou, A., Steingroever, H., Derks, K., Rouder, J.N., Morey, R.D., 2017. Bayesian inference for psychology. Part II: Example applications with JASP. Psychon. Bull. Rev. doi:10.3758/s13423-017-1323-7

